# Scanning ion-conductance microscopy for studying β-amyloid aggregate formation on living cell surface

**DOI:** 10.1101/2022.06.30.498242

**Authors:** Vasilii S. Kolmogorov, Alexander S. Erofeev, Evgeny P. Barykin, Roman V. Timoshenko, Elena V. Lopatukhina, Sergey A. Kozin, Sergey V. Salikhov, Natalia L. Klyachko, Vladimir A. Mitkevich, Christopher R.W. Edwards, Yuri E. Korchev, Alexander A. Makarov, Petr V. Gorelkin

## Abstract

Alzheimer’s disease (AD) is the most common form of dementia, a progressive neurological disorder characterized by short and long-term memory loss, including cognitive and functional impairment, which is refractory to current therapy. It is suggested that the aggregation of β-amyloid (Aβ) peptide on neuronal cell surface leads to various deviations of its vital function due to myriad pathways defined by internalization of calcium ions, apoptosis promotion, reduction of membrane potential, synaptic activity loss etc. These are associated with structural reorganizations and pathologies of the cell cytoskeleton mainly involving actin filaments and microtubules, and consequently – alterations of cell mechanical properties. Thus, the effect of amyloid oligomers on cells’ Young’s modulus has been observed in a variety of studies. However, the precise connection between the formation of amyloid aggregates on cell membranes and their effects on local mechanical properties of living cells is still unresolved. In this work, we have used correlative scanning ion-conductance microscopy (SICM) to study cell topography, Young’s modulus mapping and confocal imaging of Aβ aggregates formation on living cell surfaces with subsequent assessment of the reactive oxygen species levels inside single cells using platinum nanoelectrodes. We showed that correlative SICM technique, in conjunction with topography mapping and confocal imaging, can be used for Patch-Clamp recordings from living cells with evidently formed FAM-labeled Aβ aggregates on its surface. As we demonstrated, SICM can be successfully applied to studying cytotoxicity mechanisms of Aβ aggregates on living cell surface.

## Introduction

Neurodegenerative diseases such as Alzheimer’s disease (AD), are characterized by the formation of toxic protein aggregates that cause dysfunction and death of neuronal cells^1–4^. Extensive neurodegeneration in AD is preceded by early synaptic disorders associated with changes in the membrane ion conductivity of neuronal cells^5–7^. It has been shown that receptor-mediated toxicity and disruption of ion channel function can induce subsequent stages of the pathogenic cascade, such as pathology of tau proteins and neuroinflammation in AD^8–10^. Also, it is known that the development of neurodegenerative diseases is closely associated with dysfunction of the cytoskeleton. Oligomers of β-amyloid peptide (Aβ) cause hyperphosphorylation and aggregation of the tau protein, which leads to destabilization of microtubules. β-amyloid also mediates alterations of the actin cytoskeleton - on the one hand, due to the activation of the kinases LIMK1, p38MAPK, CAMKII, Rho/Cdc42 GTPases, and on the other hand, due to the induction of oxidative stress, leading to glutathionylation and polymerization of actin filaments^11–16^. The state of the cytoskeleton is associated with a biophysical property, Young’s modulus of the cell, which is the most important mechanism for regulating the functions of membrane proteins. Treatment with Aβ oligomers leads to a pathological change in membrane properties, but the data with regard to the direction of this change and the mechanisms involved are contradictory. Most of the results have been obtained using Atomic Force Microscopy (AFM), which exerts an adverse effect on the scanned cells due to high force applied. AFM is widely used to study the morphology and self-assembly of amyloid aggregates^17–20^, as well as for monitoring interactions between amyloid aggregates and other biological systems. However, deformation of soft cell features and other biological samples with a cantilever is a significant disadvantage of this method^21,22^.

Scanning ion-conductance microscopy (SICM) measures local mechanical properties of cells in cell culture^23,24^, making it possible to study the contributions of various molecular mechanisms to changes in membrane properties mediated by β-amyloid and potentially identify molecular targets to prevent or ameliorate this effect. Fusco et al. recently reported^25^ their observation of loss of membrane integrity in human neuroblastoma cells during incubation with toxic oligomers of α-Syn using confocal scanning microscopy. Although fluorescence-based imaging techniques are sensitive and efficient in determining cell permeability, it is still difficult to use these to obtain direct evidence for membrane disruption or morphological change in cell membranes. For a more complete understanding of the properties of such protein aggregates on cell membranes, a high-resolution surface imaging technique is required. SICM is capable of imaging cell surfaces, proteins in the membranes of living cells with nanoscale resolution and specific ion channels^26–29^. Thus, with this method it is possible to visualize the possible pore formation caused by α-Syn, with a resolution diameter of several nanometers^30^. Moreover, SICM can be used for local delivery / excitation and detection of various analytes^31,32^ as well as to study individual aggregates of beta-amyloid, causing a temporary influx of calcium through the cell membrane of neuronal cells32. Recently, SICM has been used to examine the effect of α-Syn aggregates on SH-SY5Y neuroblastoma cell membranes, a novel approach to study neurological disorders^30^. Studies were performed with cells pre-cultured with α-Syn aggregates; when using SICM, dramatic violations of cell membranes were observed. The potential role of extracellular α-Syn in pathogenesis is supported by the observation of α-Syn oligomers in the cerebrospinal fluid of patients with Parkinson’s disease^34^. In addition, a recent electrophysiological study of hippocampal neurons has shown that extracellular α-Syn has a significant effect on membraneintegrity. However, SICM visualization of beta-amyloid aggregates on live cells as well as Young’s modulus alteration and correlative confocal imaging have not yet been reported.

Thus, the nanopipette, as a key element of SICM, can be used as a nanoscale sensor. Sensors based on nanopipettes are used to determine the kinetics of formation of intracellular reactive oxygen species (ROS), the oxygen gradient and the profile of local pH values in neuronal cells and tissues^35–37^. Hence, this technique may be successfully used for studying connection of ROS generation and local mechanical properties of cells treated with β-amyloid.

In this study we have demonstrated the unique advantages of SICM for studying Aβ aggregate formation on living SH-SY5Y cell membranes, which play a key role in the development of Alzheimer’s disease. We first observed the disruption of lipid membranes in response to localized amyloid aggregate. We then measured the reduction of membrane potential by performing current clamp recordings using whole cell mode. We then successfully made intracellular ROS level measurements using nanoelectrodes and showed increased ROS level in cells induced by formed aggregates on the cell surface.

## Results and Discussion

In this work, using SICM we have revealed formation of abnormal structures on living cell surfaces after 4 hours of incubation with 10 μM Aβ_42_, (Fig.1 a-d), which is similar to α-Syn aggregates described previously^30^. The pS8-Aβ_42_ and isoD7-Aβ_42_ peptides are post-translationally modified forms of β-amyloid, which are found *in vivo* and could be significant for pathogenesis of AD^38–40^. Moreover, the formation of such structures has been accompanied by dramatic increase of Young’s modulus of the whole cell (S1). In contrast local Young’s modulus at the area of the formed structure is reduced (Fig.1-il).

**Figure 1.**
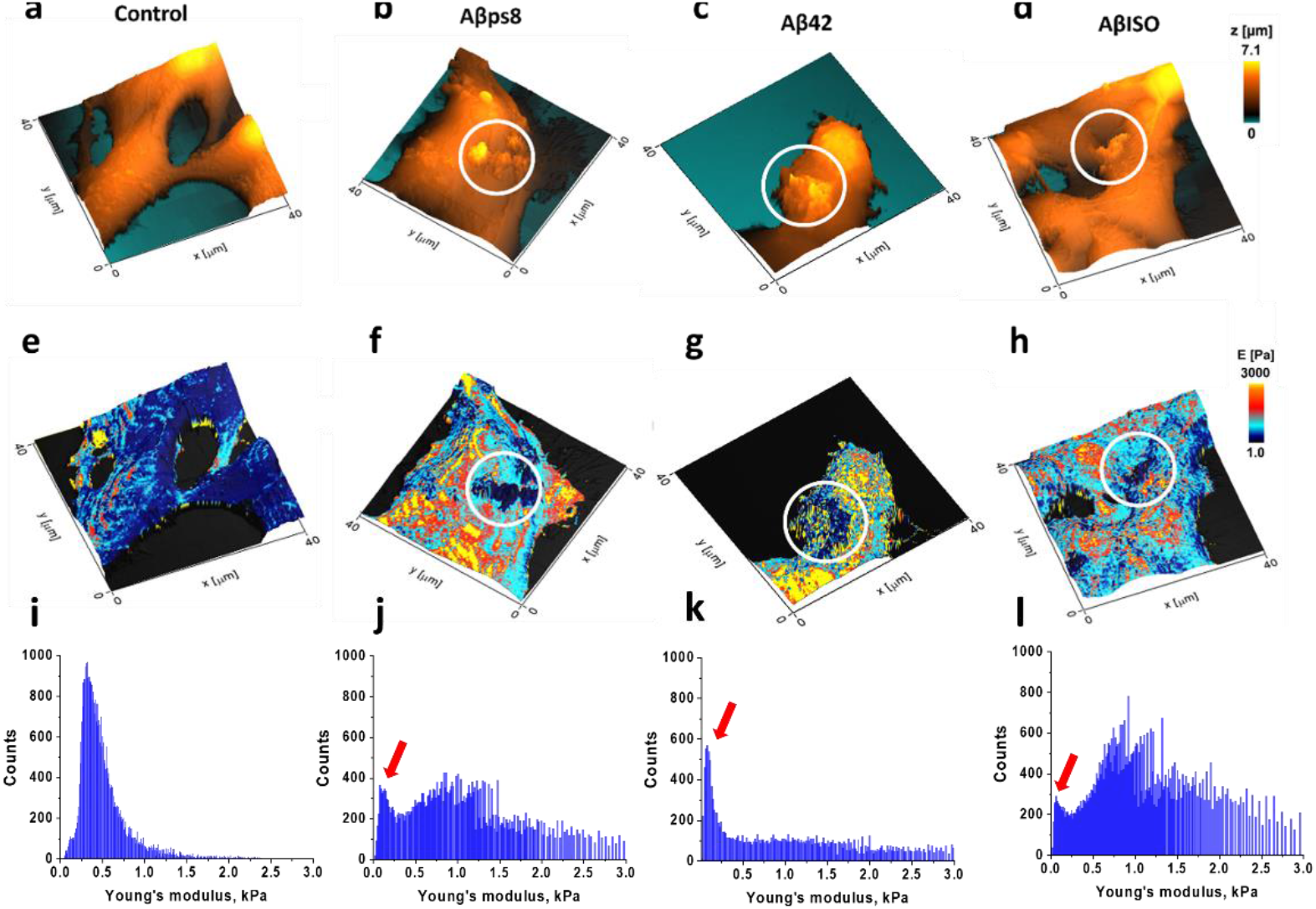
Topography (a) and Young’s modulus map (e) of SH-SY5Y control cell. Topography of SH-SY5Y cell surface after treatment with, Aβps8 (b), Aβ_42_ (c) and AβISO (d); respective Young’s modulus map (e – h); Young’s modulus distribution histogram (i) of SH-SY5Y control cell. Young’s modulus distribution histogram after treatment with, Aβps8 (j), Aβ_42_ (k) and AβISO (i); (red arrows) typical peaks on Young’s modulus distribution histogram corresponding to formed soft structures

It is known that incubation with β-amyloid oligomers leads to a significant increase in the amount of fibrillar actin, which may contribute to the cell’s Young’s modulus as measured by SICM. The disturbance of calcium homeostasis caused by Aβ_42_ leads to the activation of Rho GTPases, which are key regulators of F-actin polymerization^41^. Thus, using the SICM technique, we can confirm the changes in Young’s modulus distribution in living cells induced by Aβ_42_ and its isoforms. Using Young’s modulus as an integral characteristic of the state of the cytoskeleton in cells, we can determine which mechanisms lead to changes in the dynamics of the cytoskeleton under the action of Aβ_42_. Aβ_42_ has been reported to induce cytoskeletal modifications such as microtubule disassembly and actin polymerization leading to synaptic degeneration in neurons^13^.

These effects lead to changes in the mechanical properties of cells, which can be assessed using the SICM technique. In addition, we used fluorescently labeled Aβ_42_ (FAM-Aβ_42_) to study the structure, shape and size of Aβ aggregates on the surface of the cell membrane (Figure 2 a-c). The possibility of studying the structure of such protein aggregates was demonstrated by an example of α-synuclein when exposed to compounds based on cinnamic acid derivatives that prevent the aggregation process. It is clearly observable that β-amyloid aggregates form pore-like structures embedded in the cell membrane (Figure 2 d-e). The formed structures on Figure 2 (d-g) have 5,4 μm^2^ in area and 4,5 μm in length.

**Figure 2.**
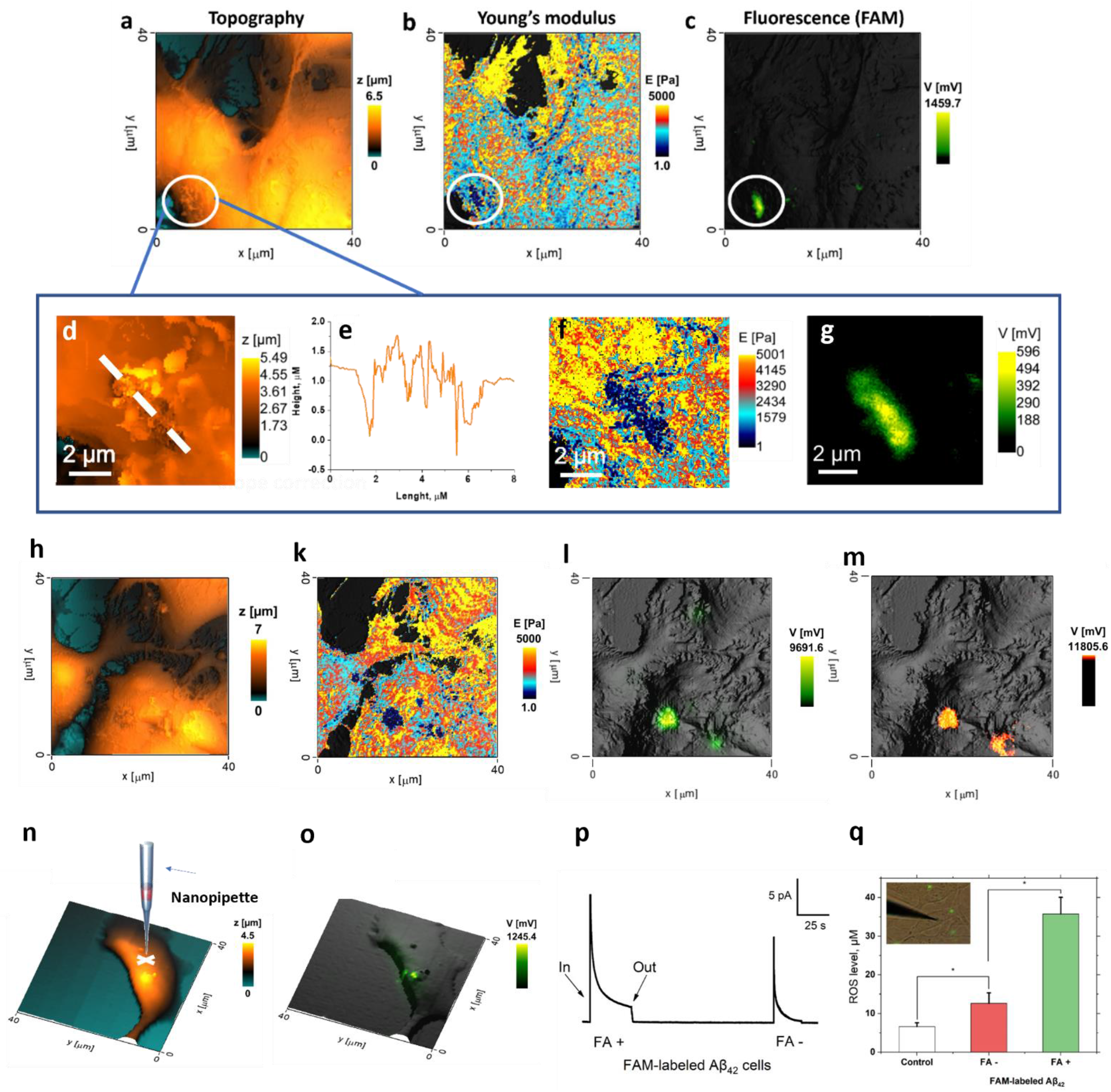
Scaling of amyloid aggregate formation on living cell membrane. Correlative imaging of topography (a), Young’s modulus (b), confocal imaging of FAM-labelled Aβ_42_; Magnified correlative images of topography (d), (e) surface profile, (f) Young’s modulus (g), confocal imaging of FAM-labelled Aβ_42_ (h-m) and DiD-lipid dye. Possible mechanism of amyloid aggregate integration to living cell membrane. Measurement (n-o) of membrane potential of control cells and cells treated by FAM-labelled Aβ_42_ via Patch-Clamp/Whole cell; (p) The difference between the current response of single SH-SY5Y cells. Cells with fluorescent aggregate were designated as «FA +», cells without fluorescent aggregate – «FA -»; (q) ROS measurements inside the SH-SY5Y cells treated by FAM-labeled Aβ_42_ for 4 h in the culture media. Cells with fluorescent aggregate were designated as «FA +», cells without fluorescent aggregate – «FA -». Untreated cells were used as a control sample. Inset on graph - merged optical and fluorescent image. The results are shown as mean, SE and (*) p < 0.05 (one-way ANOVA)

In previous works, including those performed on SH-SY5Y neuroblastoma cells, the alterations in cell stiffness were induced by treatment with oligomeric preparations of Aβ obtained with *in vitro* aggregation^12,13^. Here, the changes in Young’s modulus were observed in cells treated with fresh preparations of Aβ_42_, previously shown to contain >80% monomers^42^. Since the formation of oligomers under the specified experimental conditions is unlikely, the monomeric forms of Aβ seem to be also capable of influencing the mechanical properties of SH-SY5Y neuroblastoma cells. Also, it is evident that the interaction with the cell membrane markedly accelerates the aggregation of Aβ_42_ leading to formation of large structures on the cell surface.

We then used correlative SICM to obtain the topography (Figure 2h), Young’s modulus distribution (Figure 2k) and confocal imaging living SH-SY5Y cells simultaneously treated with FAM-Aβ_42_ (Figure 2i) and DiD lipid dye (Figure 2m). As we can see, abnormal structures on the cell topography and the local decrease in Young’s modulus at the same spot are highly co-localized with fluorescent signal of FAM-Aβ_42_ and the DiD lipid dye. We presume that high intensity DiD dye fluorescence corresponds to increased concentration of lipids on focal plane and local disruption of cell membrane. Detergent-like properties of amyloid aggregates were reported previously^43^, but commonly have been shown using artificial cell membrane models such as liposomes or lipid bilayers, and not on living cells^44–46.^

Correlative SICM allows us to perform Patch-Clamp recordings on cells where membrane-bound β-amyloid aggregates were detected with topography mapping and confocal imaging (Figure 2). In such cells, we observed lower membrane potential (−30 mV) compared to control cells (−50 mV). Reduced membrane potential (S2) is possibly connected to metabolic alteration like ROS generation and mitochondrial disfunction^47,48^ leading to ionic homeostasis impairment.

Further, we used Pt nanoelectrodes for intracellular ROS measurements. In cells with membrane-bound FAM-Aβ_42_ aggregates we noticed a significant increase (Figure 3) in ROS related to the ability of Aβ_42_ to disrupt the mitochondria and induce oxidative stress, as well as to Aβ-induced calcium influx^48^. Notably, ROS levels measurements in cells treated with FAM-Aβ_42_ without aggregates on the membrane, demonstrated induced oxidative stress but to a lower extent than cells with membrane-bound aggregates. Higher cytotoxicity of amyloid aggregates than free oligomers is clear, a matter which was unresolved in previous studies.

## Conclusions

In this work, correlative SICM was successfully used as a multimodal technique for studying β-amyloid aggregate formation on the surface of living SH-SY5Y cells. We have shown that there is an increase of Young’s modulus of the whole cell during treatment with β-amyloid and its modified forms and the formation of abnormal structures characterized by locally reduced Young’s modulus. Simultaneous confocal imaging allowed us to detect pore-like formation of β-amyloid aggregates followed by local membrane disruption leading to a reduction in membrane potential. Pt nanoelectrode application demonstrated higher toxicity and oxidative stress induction by membrane-bound aggregates in comparison to soluble oligomers of β-amyloid.

## Methods and Methods

### Amyloid peptides preparation

Synthetic peptides (purity > 98% checked by RP-HPLC) Aβ_42_(DAEFRHDSGYEVHHQKLVFFAEDVGSNKGAIIGLMVGGVVIA), fluoresceine amidite-Aβ_42_ (FAM-Aβ_42_), pS8-Aβ_42_ and isoD7-Aβ_42_ were purchased from Biopeptide Co., LLC (San Diego, CA, USA). Peptides were monomerized as described previously^49^. A fresh 2.5 mM solution of Aβ was prepared by adding 20 μl of 100% cell culture grade anhydrous dimethyl sulfoxide (DMSO) (“Sigma”) to 0.224 mg of peptide, followed by incubation for 1 h at room temperature to completely dissolve the peptide.

### Cell preparation

SH-SY5Y neuroblastoma cells (ATCC, USA) were cultivated according to a standard protocol in Dulbecco’s Modified Eagle’s Medium (DMEM/F-12, Gibco, USA) with 1% L-glutamine and 4-(2-hydroxyethyl)-1-piperazine-ethanesulfonic acid (HEPES) (15 mM, Fisher) supplemented with fetal bovine serum (10% (v/v), Gibco, USA) and penicillin–streptomycin (1%) (Research Product International, Mt Prospect, IL). TrypLE (0.25%, Fisher) was used to detach cells from the culture plate and fresh culturing medium was added every 3–4 d.

SH-SY5Y (3 × 10^5^) cells were seeded in 35 mm Petri dishes and treated the next day with either Aβ_42_, FAM-Aβ_42_, pS8-Aβ_42_ or isoD7-Aβ_42_. The amyloid peptides were dissolved in DMSO as described above and diluted in serum-free culture medium. The final concentration of the peptides in the culture medium was 10 μM with 4 h of incubation time. Cells treated with an equivalent amount of DMSO in culture medium without amyloid peptides were used as a control, which was performed at the beginning and at the end of the experiment. Before imaging, cells were washed three times with Hanks’ Balanced Salt solution to remove the growth media and traces of FAM-labelled Aβ_42_. For confocal imaging of cell membranes, Vibrant DiD-lipid dye (Thermofisher, USA) was used at a final concentration of 1 μM.

### Scanning Ion-Conductance Microscopy (SICM)

Scanning procedure was performed using SICM manufactured by ICAPPIC (ICAPPIC Ltd, United Kingdom). Nanopipettes with a typical radius 40-50 nm were drawn from borosilicate capillaries (1.0 mm, OD; 0.5 mm, ID; Sutter Instruments, USA) using a Laser Puller P-2000 (Sutter Instruments, USA).

Cell topography and QNM were performed using the non-contact hopping mode with an adaptive resolution with maps of 40 × 40 μm, with image resolution of 256 × 256 pixels. The approach rate during imaging was set at 100 μm s-1. For all SICM measurements, a three-set point mode was used. A non-contact topographic image was obtained at an ion current decrease of 0.5% and a further two images were obtained at an ion current decrease of 1% and 2% corresponding to membrane deformations produced by intrinsic force at each setpoint. Correlative scanning ion-conductance microscopy and confocal microscopy were performed using SICM manufactured by ICAPPIC (ICAPPIC Limited, United Kingdom) equipped with a confocal module (ICAPPIC Limited, United Kingdom). l. For further correlative imaging, a 25.3 mW laser beam with a wavelength of 635 nm was focused on the nanopipette tip so that the focal plane of the laser beam and nanopipette tip were in the same place.

In this work, Patch-Clamp recordings were performed using SICM by ICAPPIC. Patch pipettes were made from borosilicate glass 1.0 mm OD and 0.5 ID (Sutter Instruments, USA) with typical electrical resistance of 10-20 MOhms using the following two-step program - HEAT1 310, FIL1 3, VEL1 30, DEL1 150, PUL1 0, HEAT2 300, FIL2 4, VEL2 25, DEL2 150, PUL2 200.

### Patch-Clamp recordings

Submicron pipettes with a typical radius 0.5-1.0 μm were used for scanning cell topography with correlative confocal imaging by SICM ICAPPIC to find SH-SY5Y cells with formed aggregates of FAM-Aβ_42_ on the cell membrane (Fig.2 n, o). A giga-ohm seal was then made by applying light negative pressure near the cell membrane. Whole-cell mode through applying sharp negative pressure has been performed. After that membrane potential was measured at current zero mode.

### Intracellular ROS level measurements

The total ROS concentration was determined by the amperometric method using Pt-nanoelectrodes. Commercially available disk-shaped carbon nanoelectrodes isolated in quartz (ICAPPIC Limited, UK) with diameters 60–100 nm were used to prepare Pt nanoelectrodes. Firstly, the carbon surface was etched in a 0.1 M NaOH, 10 mM KCl solution during 40 cycles of 10seconds (from 0 to +2200 mV) to create nanocavities. Further electrochemical deposition of platinum in nanocavities was achieved by cycling from 0 to 800 mV with a scan rate of 200 mV/s for 4 to 5 cycles in 2 mM H2PtCl6 solution in 0.1 M hydrochloric acid. Cyclic voltammetry from −800 to 800 mV with a scan rate of 400 mV/s in a 1 mM solution of ferrocene in methanol in PBS was used to control the electrode surface at all stages of fabrication. Prior to the measurements, each platinum nanoelectrode was calibrated using a series of standard H2O2 solutions at a potential of +800 mV. Preparation of Pt-nanoelectrodes has been described in detail elsewhere.

A nanoelectrode penetrated the cells and measured the oxidation current of hydrogen peroxide. On average, about 8 cells were measured by 2–4 Pt electrodes in independent Petri dishes. To confirm that fibril-like formations on SH-SY5Y cell surface are indeed amyloid aggregates, FAM-Aβ_42_ was used in similar conditions as Aβ_42_ which are described earlier.

## Author Contributions

The study was performed employing unique scientific facility “Scanning ion-conductance microscope with a confocal module” (registration number 2512530) and was financially supported by the Ministry of Education and Science of the Russian Federation, Agreement No. 075-15-2022-264. Yu. Korchev acknowledges the support from the World Premier International Research Center Initiative (WPI) from the MEXT of Japan.

