## Supplementary material for "Scanning ion-conductance microscopy for studying β-amyloid aggregate formation on living cell surface": Supplemetary

### Electronic Supplementary Information

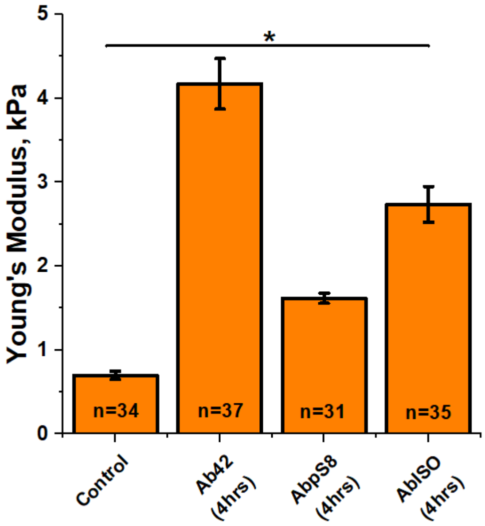


Figure 1S. Mean value of Young's modulus of control cells, cells treated by 10 μM A$\beta_{42}$, pS8-A$\beta_{42}$ and isoD7-A$\beta_{42}$ (n – amount of measured cells, * $p \leq 0.05$ One-Way ANOVA)

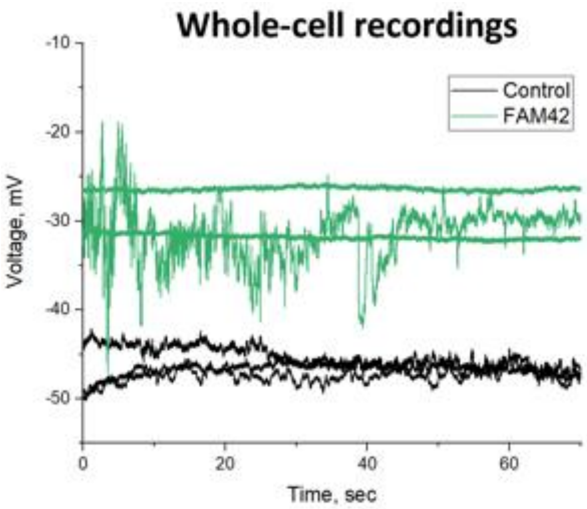


Figure 2S. Membrane potential recordings of control cells (black line) and cells with FAM-A$\beta_{42}$ aggregates
